# Conflict and cooperation in eukaryogenesis: implications for the timing of endosymbiosis and the evolution of sex

**DOI:** 10.1101/023077

**Authors:** Arunas L. Radzvilavicius, Neil W. Blackstone

## Abstract

The complex eukaryotic cell is a result of an ancient endosymbiosis and one of the major evolutionary transitions. The timing of key eukaryotic innovations relative to the acquisition of mitochondria remains subject to considerable debate, yet the evolutionary process itself might constrain the order of these events. Endosymbiosis entailed levels-of-selection conflicts, and mechanisms of conflict mediation had to evolve for eukaryogenesis to proceed. The initial mechanisms of conflict mediation were based on the pathways inherited from prokaryotic symbionts and led to metabolic homeostasis in the eukaryotic cell, while later mechanisms (e.g., mitochondrial gene transfer) contributed to the expansion of the eukaryotic genome. Perhaps the greatest opportunity for conflict arose with the emergence of sex involving whole-cell fusion. While early evolution of cell fusion may have affected symbiont acquisition, sex together with the competitive symbiont behaviour would have destabilized the emerging higher-level unit. Cytoplasmic mixing, on the other hand, would have been beneficial for selfish endosymbionts, capable of using their own metabolism to manipulate the life history of the host. Given the results of our mathematical modelling, we argue that sex represents a rather late proto-eukaryotic innovation, allowing for the growth of the chimeric nucleus and contributing to the successful completion of the evolutionary transition.

## Introduction

The endosymbiotic theory of the origin of the complex cell has fascinated biologists for over a century (e.g., [1]). Molecular phylogenetics has identified the symbionts [2], narrowed the possible candidates for the host [3], and with chronological calibration provided estimates for the timing of eukaryotic origins [4]. Comparative genomics suggest that the common ancestor of all extant eukaryotes exhibited virtually all of the derived characters of the group [5]. What remains difficult to elucidate is the actual process of eukaryogenesis. Because this process occurred in the eukaryotic stem group, examining modern taxa remains uninformative in this context (Figure 1). Several outstanding questions thus remain [6]; in particular, how and when were mitochondria acquired relative to the defining features of eukaryotes? To the extent that eukaryotic features can be considered sequelae to the endosymbiosis, progress can be made in answering these questions. For instance, the definitive shared derived character of eukaryotes, the nucleus, can be interpreted as a barrier between transcription and translation that evolved as slow-splicing, proto-mitochondrial introns became abundant in the proto-nuclear genome [7, 8].

**Figure 1.**
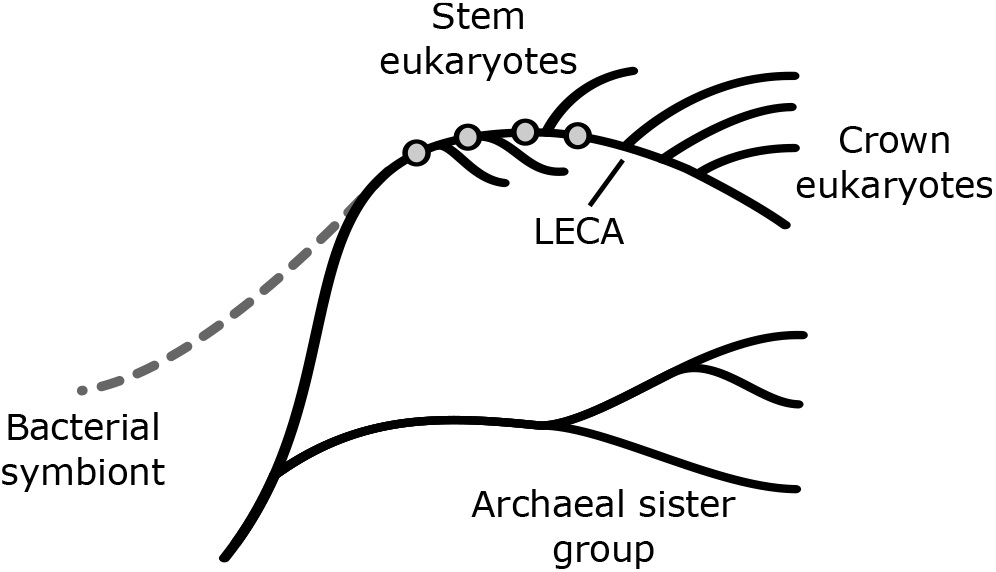
Schematic tree representation of the endosymbiotic event leading to the origin of eukaryotes. The last eukaryotic common ancestor (LECA) exhibited virtually all key eukaryotic features, including sex. The relationship between the acquisition of the defining features of eukaryotes and the timing of the endosymbiosis, however, remains a subject of controversy. Any features of eukaryotes that can be considered sequelae to the endosymbiosis must have evolved subsequent to the acquisition of bacterial endosymbionts.

A broader understanding of the consequences of the endosymbiosis can be obtained by an evolutionary analysis of conflict and cooperation. There is a general agreement that major transitions in biological complexity are driven by cooperative interactions between initially independent units at the lower level of organization [9]. A stable community of cooperators under certain conditions can be transformed into a new unit of selection, while individuality at the lower level is lost in whole or part. Due to the strong individual selection, however, conflicts between levels of organisation are hardly avoidable; competition and selfish behaviour of the lower-level units destabilizes and may eventually destroy the emerging higher level of individuality. Mechanisms repeatedly mediating such conflicts and preserving stable community interactions are therefore necessary [10]. In this regard, the origin of eukaryotes was likely similar to other major transitions.

The initial relationship between the symbiotic partners was likely very different from modern cells, although the exact nature of the metabolic association leading to the proto-eukaryotic endosymbiosis is unclear [11]. Regardless of the initial reasons bringing the two prokaryotic species together, the stability of the symbiont community residing inside of the archaeal host was threatened by the competition between individual proto-mitochondria. Under some circumstances, uncontrolled selfish replication and high reactive oxygen species (ROS) emissions could result in killing the host with symbionts returning back to the free-living state or spreading throughout the population via repeated host cell fusions and divisions [12]. The central role of modern mitochondria in executing the apoptotic cell death programme, for example, can be interpreted as a consequence of such conflict [13 – 16].

Gene transfer to the nucleus and subsequent reduction of the proto-mitochondrial genome serves as one way to mediate selfish conflicts. Genome transfer restricts the mutational space and decreases heritable variation between the lower level units, at the same time centralising the control of many essential proto-mitochondrial functions at the emerging chimeric nucleus [17]. Genomes of modern mitochondria are never capable of controlling their own replication or energy allocation, but selection on the lower level would initially oppose the loss of such genes. Additionally, genome transfer required numerous evolutionary innovations, such as novel mitochondrial protein import machinery, suggesting that gene transfer did not play a significant role in mediating conflicts during the early stages of eukaryogenesis.

Alternative mechanisms of conflict mediation may have existed from the very beginning of the endosymbiotic association. Indeed, mitochondrial evolution can be conceptualized as cycles of cooperation and conflict. With each cycle, new conflicts arise and must be mediated [10]. A closer look at the early evolutionary history of mitochondria suggests that initial mechanisms alleviating conflicts were likely driven by individual selection on proto-mitochondria in the new environment, that is, the emerging cytosol where tight bioenergetic membrane coupling, selfish energy allocation and rapid replication could lead to damage and eventual demise of the lower-level units. These very same mechanisms became the basis of the novel signalling pathways and means of communication between the levels of selection in the emerging eukaryotic communities.

Sex is among the numerous features of complex cell that originated during the evolutionary transition towards eukaryotes [18]. The evolution of sex has been traditionally analysed in terms of costs and benefits of chromosome segregation and recombination, paying little attention to the historical circumstances such as the process of eukaryogenesis. Cell fusion, for example, may be viewed as a result of mitochondrial manipulations allowing endosymbionts to spread through the population without the rigors of a free-living stage, at the same time allowing for the reciprocal recombination stabilizing the newly emerging nuclear genome and restoring favourable conditions for growth and replication [16]. Mixing of mitochondrial populations from different lineages has deleterious effects as demonstrated by nearly universal uniparental inheritance of organellar DNA in modern eukaryotes. With similar selective forces operating during the eukaryogenesis, cell fusion and cytoplasmic mixing would have been costly for the emerging higher-level unit, potentially disrupting the evolutionary transition.

The conflict and cooperation that accompanied the levels-of-selection transition can thus provide insight into the evolution of eukaryotic features in general and sex in particular relative to the timing of the mitochondrial endosymbiosis. Given this background, a number of canonical features of eukaryotes will first be reinterpreted as mechanisms of conflict mediation. Second, a simple model will be used to examine the effects of sex on the fitness of the higher-level unit. We will explicitly consider the spread of alleles coding for the cell-cell fusion in the initially asexual population. Frequent mixing of cytoplasmic contents can significantly undermine the fitness on the higher level of selection given strong competition on the lower level. Selfish conflict therefore has to be suppressed before sex can evolve. Our analysis suggests that many integral features of eukaryotes must necessarily have evolved subsequent to the seminal endosymbiotic event and constrains the extent to which eukaryogenesis may have preceded endosymbiosis.

### Individual selection within the emerging cytosol

A great number of hypotheses exists for the initial symbiotic association between bacterial ancestors of mitochondria and their archaeal partners [11]. Independent of the metabolites involved, the initial association eventually led to the endosymbiosis with multiple proteobacteria residing within the host’s cytoplasm. While in modern eukaryotes the competition between mitochondria is largely suppressed, during the earliest stages of eukaryogenesis selection among symbionts was still present, and the new environment might have offered a selective advantage such as the larger area of contact ensuring more efficient exchange of metabolites. The stable intracellular community lifestyle, however, required some radical changes and evolutionary innovations.

The metabolic state of a cell depends on a series of electron couples with environmental sources and sinks [19]. It is therefore the environment that ultimately sets the capacity for growth and replication. For proteobacterial endosymbionts the immediate environment was the proto-eukaryotic cytosol, which itself was an emergent feature of the community. To a large extent the metabolic state of an individual endosymbiont was being controlled by the whole proto-mitochondrial collective as well as the host. In cases where such community control is harmful for some of the proto-mitochondria, individual selection would result in new adaptations increasing the chances of survival.

Consider the internal environment of the emerging eukaryote, supplying proto-mitochondria with an adequate amount of substrate, but limiting their growth and proliferation. Due to low metabolic demand the endosymbionts would likely convert most of their ADP into ATP and experience high levels of NADH relative to NAD^+^. The resting state 4 metabolism [20] would ensue, with proto-mitochondrial electron transport chains becoming highly reduced. Highly reduced electron carriers freely contribute electrons to molecular oxygen thus producing dangerously large amounts of ROS (Figure 2) [21]. ROS would likely damage not only the symbiont genomes, but also its proteins and lipids, possibly leading to eventual demise.

**Figure 2.**
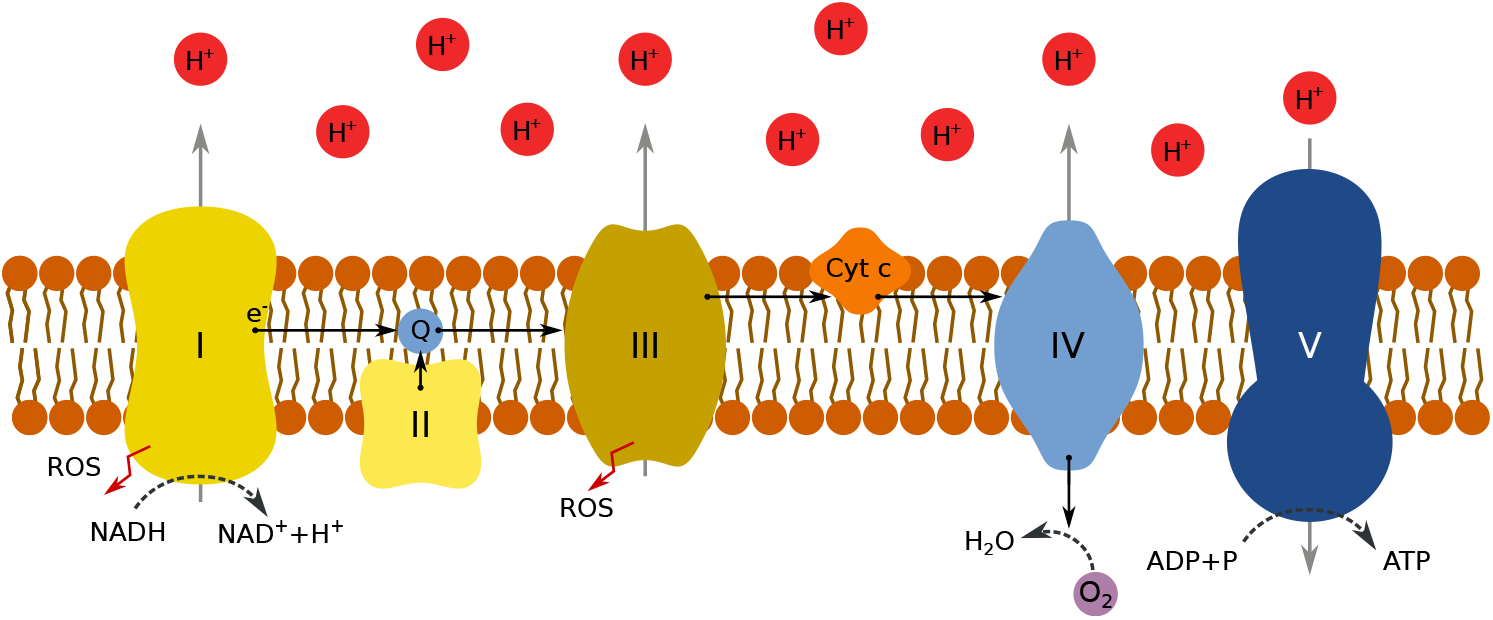
Schematic representation of the mitochondrial electron transport chain. Substrate is oxidized and electrons follow a path (black arrows) through the electron carriers of complexes I – IV to molecular oxygen. Protons are extruded (gray arrows) and return to the mitochondrial matrix via ATP synthase (V). In state 3, substrate supply is adequate, and metabolic demand is high. The redox state of the electron carriers, the proton gradient, ROS, and ADP levels are intermediate. Phosphorylation rates are maximal. In state 4, substrate supply is adequate, but metabolic demand is weak. ADP relative to ATP is minimal, the proton gradient is maximal, electron carriers are highly reduced, and ROS formation is maximal.

**Figure 3.**
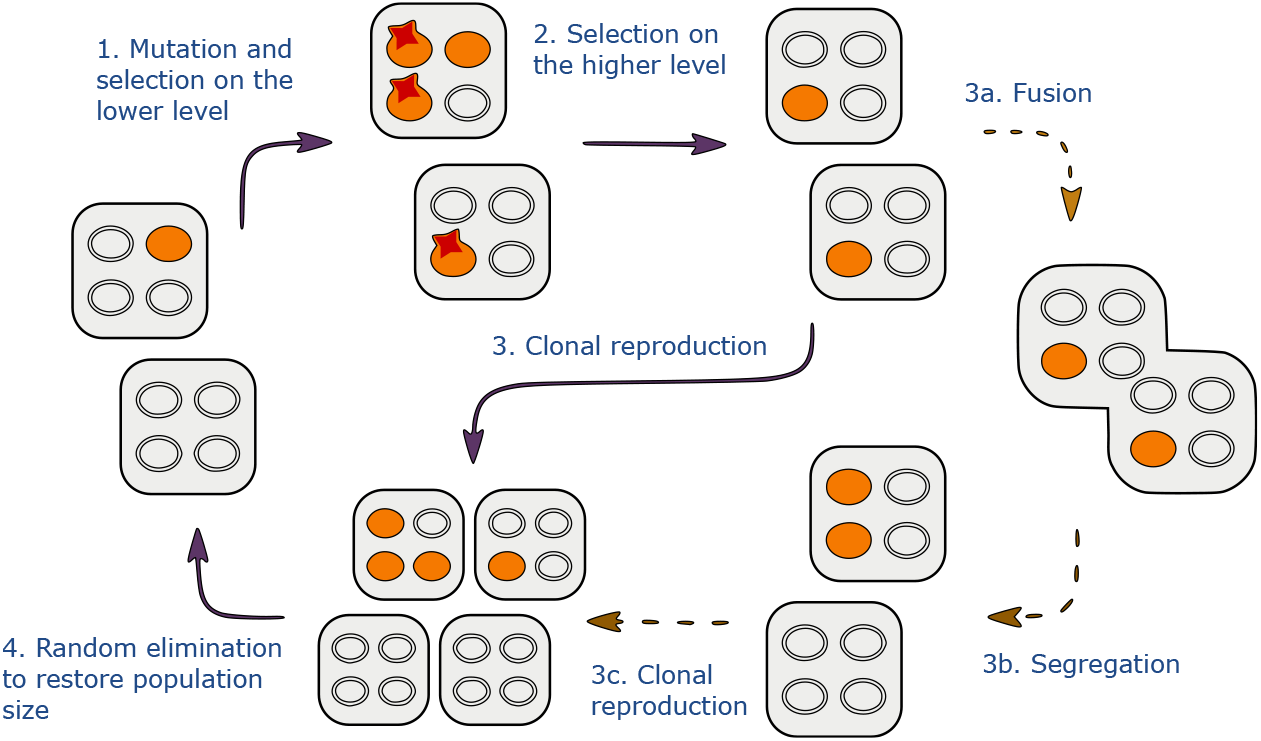
Schematic representation of the model life cycle. Unicellular organisms containing *M* endosymbionts undergo steps 1-4 as described in the main text. Shaded symbionts represent selfish mutants, while wild-type proto-mitochondria are left blank. Depending on the allele at the nuclear locus *A/a*, an organism can reproduce clonally or temporarily fuse with a random partner mixing the cytoplasmic contents.

ROS-damaged proto-mitochondria would disappear more rapidly if the cytosol were able to selectively digest damaged endosymbionts in the process analogous to modern mitophagy. Such cytosolic consumption would be of particular importance if the host were not able to perform oxidative phosphorylation, and had to rely on substrate-level-phosphorylation with high substrate demands [22], for example, due to increased size and low surface area-to-volume ratio. Furthermore, the degradation of damaged endosymbionts would be necessary to prevent further damage to the rest of the community by both ROS and possible toxic mitochondrial constituents.

Under such conditions, individual selection favours symbionts that in one way or another manage to avoid state 4 metabolism and oxidative damage. Indeed, much of the early evolution of mitochondria may have led to the establishment of metabolic homeostasis in this novel, challenging environment. For example, mutant mitochondria with weak coupling between electron transport and ATP production, thus wasting part of the energy in the form of heat, would be able to persist in state 3 metabolism indefinitely, despite low metabolic demand dictated by the cytosol. Similarly, free-living bacteria employ the energy spilling mechanism [23], illustrating the dangers of metabolic state 4, which could potentially be even greater within the foreign cytoplasm. Under certain conditions, proto-mitochondrial metabolic inefficiency can therefore be selected for, despite the fact that it results in lower rates of growth and proliferation during the favourable times.

While proto-mitochondrial DNA mutations lowering the efficiency of membrane coupling offered some early advantages under stressful conditions, symbionts with active control of their own homeostasis would eventually evolve. Such proto-mitochondria would retain high efficiency of ATP production when metabolic demand is high, while still being able to avoid overproduction of ROS and cytosolic digestion under less favourable environmental conditions. There are a number of ways this can be achieved, and much of the signalling in a modern cell relates to these very issues [24 – 26]. Nevertheless, some modern mechanisms were likely more difficult to evolve than others. Likely, initial mechanisms were co-opted from features already present in bacteria.

For instance, consider calcium signalling in this context [27]. Low intracellular calcium concentration relative to the external environment is a shared feature of all life, most likely because the first living cells emerged within a low-calcium environment. As a consequence, calcium ion influx from either the external environment or intracellular stores can be easily used as a primitive but highly versatile signal. In modern eukaryotic cells, mitochondria alongside the endoplasmic reticulum (ER) appear to be central in calcium signalling pathways.

The physical association of mitochondria with the calcium transporters (e.g. inositol-triphosphate receptors in the ER) has an elegant evolutionary explanation in the conflictual stages of early eukaryogenesis. The bacterial ancestors of mitochondria could have used the regulated Ca^2+^ uptake to depolarize their external membranes and avoid entering metabolic state 4. Furthermore, the subsequent extrusion of calcium ions by ATP-dependent pumps would have provided an additional source of metabolic demand, lowering the ATP/ADP ratio and thus sustaining the type 3 metabolism even in the conditions of low metabolic demand. Due to the low overall intracellular calcium concentrations only proto-mitochondrial mutants capable of establishing physical associations with the host’s membranes and calcium influx channels would have been able to maintain their tight membrane coupling and still avoid the dangers of ROS overproduction. Lower rates of damage and digestion within the cytosol would have therefore driven the evolution of heritable mechanisms ensuring the efficiency of proto-mitochondrial calcium uptake and eventually resulting in structures like mitochondria-associated endoplasmic reticulum membranes (MAM) [27].

Mechanisms of conflict resolution in one form or another must be present from the very beginning of the transition and therefore relevant pathways have to be recruited or co-opted from the ones existing on the lower level. Primitive calcium signalling employed to depolarize proto-mitochondrial membranes early in eukaryogenesis could be one such example. Indeed, multiple components of eukaryotic calcium signalling pathways derive from bacterial ancestors [28, 29] including recently identified homologue of mitochondrial calcium uniporter in several bacterial species [30].

### Adenine nucleotide translocase

The evolutionary transition towards a biological unit of higher complexity involves the emergence of selection and individuality on the community level, which requires communication across the levels of organisation. While early exchange of nutrients and ions initiated the transition and led to the derivation of essential features of modern eukaryotes, the real revolution arguably came with the origin of adenine nucleotide translocase (ANT).

ANT is at the centre of the modern cooperative interaction between mitochondria and the rest of the eukaryotic cell as it exports mitochondrial ATP in exchange for ADP from the cytosol. ANT belongs to a family of mitochondrial transporters of similar structure together with membrane uncoupler proteins and inorganic phosphate carriers. While it is clear that mitochondrial transporter protein family has no homologues in prokaryotes and evolved at some point during the eukaryogenesis [31], the selective forces behind the emergence of ANT are still debatable. One view posits that ANT was encoded by the host’s genome, “inserted” to exploit endosymbionts, and therefore was driven by selection on the host.

From the levels-of-selection viewpoint, the export of proto-mitochondrial ATP could be interpreted as yet another mechanism of sustaining state 3 metabolism under the conditions of low metabolic demand, when there is a shortage of ADP in the mitochondrial matrix and an overabundance of NADH. “Selfish” proto-mitochondria without ANT therefore may grow and reproduce faster under favourable environmental conditions, but would start overproducing ROS and die faster under the low metabolic demand. ANT may therefore be a consequence of individual selection between endosymbionts within the shared cytosol, since ANT-less proto-mitochondria would be more likely to suffer from oxidative damage and experience higher rates of degradation [24, 25]. If ANT evolved as a proto-mitochondrial innovation, selection would favour an increase in the frequency of ANT-containing endosymbionts. ANT could spread to the proto-nuclear genome upon release from dying endosymbionts and colonize other proto-eukaryotes via lateral gene transfer. A similar but phylogenetically unrelated case of the rickettsial nucleotide transporter illustrates that it is indeed possible for endosymbionts to gain relevant genes.

The structural and dynamical features of the ANT itself can assist in elucidating the conditions under which the mitochondrial carriers first arose. After synthesis in the cytosol, the import of ANT building blocks and their assembly in the inner membrane is dependent on mitochondrial translocases of inner and outer membranes [32]—eukaryotic innovations evolving together with the transfer of mitochondrial genes to the nucleus [33]. The traditional view of ATP/ADP transporter as the host’s innovation would therefore put the origin of ANT very late in the transition, i.e. after many endosymbiont genes have already been transferred to the nucleus and the mitochondrial protein import machinery had been established, which itself required relaxed selection and conflict mediation on the lower level.

It is also important to note that depending on the ATP/ADP gradient and the membrane potential ANT can transport nucleotides in either direction (again, similar to the rickettsial transporter) [34]. Under the conditions of high metabolic demand and the shortage of substrate, it would favour the import of cytosolic ATP (the product of fast cytosolic substrate-level phosphorylation), potentially harmful to the host. On the other hand, selection on individual proto-mitochondria would favour ANT under the conditions of both state 2 and state 4 metabolism, generating metabolic demand when the risk of ROS overproduction is high, and importing ATP when the proton-motive force across the membrane is too low.

Unlike membrane uncoupling or inefficient OXPHOS, however, the export of ATP had tremendous consequences for the community as a whole; not only did it diminish the oxidative damage of individual endosymbionts, but also provided a way to stabilize the host energetically, allowing it to grow, replicate and migrate thus possibly restoring favourable conditions for the whole collective. ATP is therefore not only the energetic currency of the eukaryotic cell, but also a language of ancient communities of cells within cells, capable of signalling metabolic state and exposing the proto–mitochondrial cheaters. While ANT is a feature that clearly evolved subsequent to endosymbiosis, its evolution triggered a number of sequelae, e.g., phosphorylation cascades, without which the eukaryotic cell would be unrecognizable.

### Genome fusion and loss

The nuclear-mitochondrial arrangement of genes and genomes represents perhaps one of the most consequential features of eukaryotes [35]. Formation of the chimeric nuclear genome was likely initiated at the same time as the endosymbiosis, while genome reduction in endosymbionts required both initial conflict mediation and later innovation (e.g., the mitochondrial protein import apparatus). The remnant genome of mitochondria allows rapid changes in gene activity in response to changing environmental conditions [19]. This can be regarded as the final chapter in the evolution of metabolic homeostasis, with the same underlying evolutionary rationale. Only mitochondria that retained components to maintain sufficient redox control of gene activity were able to maintain metabolic homeostasis, and gained a fitness advantage.

### The origin of sexual cell fusion

Despite numerous apparent disadvantages, such as breaking up co-adapted gene combinations or the need to find a partner, sex is nearly universal among the eukaryotic species. It can be shown that sex uncovers the hidden genetic variation, allows selection to “see” individual genes and gene combinations, and thus increases the efficacy of selection—but only in finite populations subject to genetic drift [36, 37]. Whole cell fusion and reciprocal sex are virtually unknown in bacteria that tend to share genetic information exclusively through lateral gene transfer, although some archaeans may exhibit more eukaryote-like mating [38]. Comparative studies of modern eukaryotes suggest that seemingly fully developed sexual life cycles were very likely present in the common ancestor of all eukaryotes [18].

Sexual life cycles suggest several important connections to mitochondrial endosymbiosis. First, the fluidity of membranes that permits whole-cell fusion might have also facilitated symbiont acquisition. Second, whole cell fusion can permit exchange of intracellular symbionts without the attendant rigors of a free-living stage of the life cycle. Additionally, Lane [39] suggested that the evolution of sex could have been driven by high mutation rates and genome instability during the phase of intense bombardment of nuclear genome by mitochondrial genes and introns, when selection on the new eukaryotic level was strong and the number of linear chromosomes was not constant.

While sex might facilitate symbiont acquisition, it also raises the spectre of symbiont mixing. Strong selection must operate against mitochondrial mixing, as demonstrated by repeated evolution of uniparental inheritance of cytoplasmic DNA [40]. Uniparental inheritance increases mitochondrial fitness variation between cells and ensures low levels of heteroplasmy [41, 42]. Similar selective pressures could have operated during the origin of eukaryotes, and therefore the effect of sex on both levels of individuality has to be considered. Sexual reproduction could represent another conflict between the levels, allowing mutant mitochondria to spread horizontally and lowering genetic variation between hosts. On one hand, sex thus would seem to simplify questions of “how” symbionts were taken up, but on the other, an initial sexual life cycle raises questions about whether a stable endosymbiosis could evolve. How challenging are the conflicts associated with symbiosis and a sexual life cycle?

### Simulation model for the origin of sexual fusion

To investigate the conditions under which sexual fusion between emerging higher-level units could be expected to evolve, we developed a mathematical model based on a finite population of haploid unicellular individuals and inspired by [41] and in part by [43]. We consider an asexual ancestral population fully dependent on the endosymbiont metabolism, and, unlike previous work, explicitly analyse the evolution of modifiers inducing sexual mixing of the cytoplasms, but ignoring the benefits associated with recombination and genome repair.

Each organism contains a fixed number of proto-mitochondrial symbionts *M*, which could be in one of two states, cooperative or selfish mutant. The model assumes that the fitness of the higher-level unit depends on the number of defectors *m*, and can be approximated as a concave function 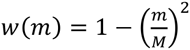.

We argue that due to the high number of symbionts within a single higher-level unit, small *m* has only a minor effect on the fitness of the host cell. As the number of mutants approaches *M*, fitness decreases more rapidly, illustrating the dependence of the higher-level unit on cooperative endosymbionts. Indeed, in many modern mitochondrial diseases high, above-threshold levels of heteroplasmy are needed to cause significant phenotypic changes [44]. As the initial fitness association between the proto-eukaryotic symbionts is not known, in the *Supplementary Material* we analyse the case of a linear fitness function 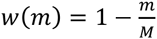 and an unlikely case of a rapidly declining convex fitness association 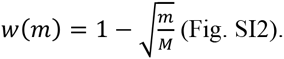 (Fig. SI2).

The life cycle is modelled as a sequence of discrete events and is illustrated in Figure 1. Selection on the higher level (step 2) is countered by rare mutations and selection between endosymbionts (step 1), with selfish mutants having a relative replication advantage 1+*k*. As a simplifying assumption, the number of individuals within the higher-level unit as well as within the population is kept constant. Higher-level units reproduce clonally, by first duplicating their contents and then randomly assigning the symbionts to the two daughter cells (step 3). Alternatively, before the clonal reproduction a cell can fuse with a random partner thus mixing the cytoplasmic contents (3a), followed by meiosis-like fission and random partitioning of endosymbionts (3b). The segregation of mutant and wild type symbionts between the two daughter cells is modelled as simple sampling without replacement, following hypergeometric probability distribution; mutation events are binomial with probability μ, while selection is modelled as the fitness-weighted sampling with replacement. The mode of reproduction (clonal or fusion) can be controlled by a single locus within the chimeric genome *A*/*a*.

### The cost of cytoplasmic mixing

To investigate the long-term effect of cytoplasmic mixing on the population fitness we first run the simulations with fixed fusion rates, independent of the allele at the fusion locus *A/a*. Since proto-mitochondrial mutations are modelled as rare events, mutants proliferate mostly due to higher replication rate within the cytosol. As we demonstrate in Figure 4, in populations that are mainly clonal selection on the higher level is efficient enough at eliminating selfish symbionts to ensure nearly complete cooperation and high community fitness. Selfish endosymbionts are mostly confined within distinct host lineages keeping the rest of the population virtually mutant-free (Figure 5a, b). This ensures high variation between higher-level units, facilitates selection and therefore results in high mean population fitness.

**Figure 4.**
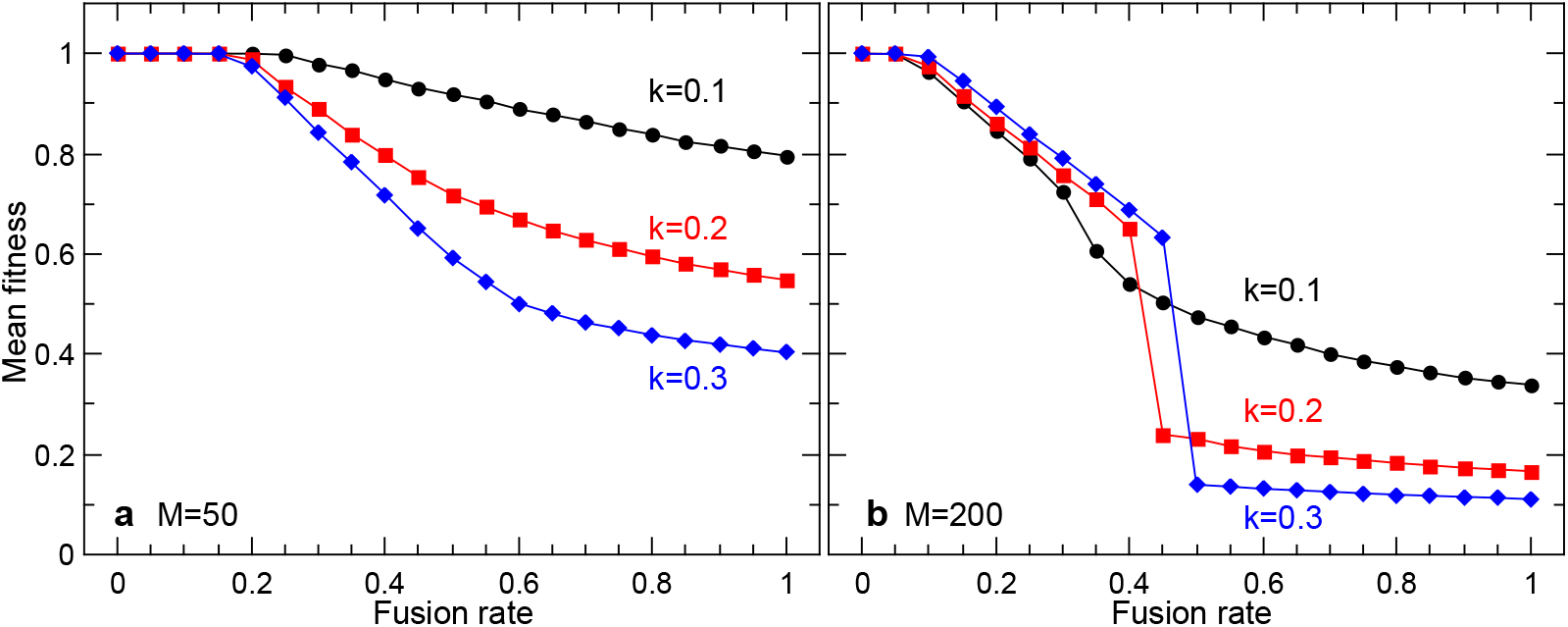
Mean population fitness as a function of the fusion rate for two cell sizes, *M*=50 (a) and *M*=200 (b). Three curves correspond to three values of mutant reproductive advantage *k*. Due to the horizontal spread of mutants and reduced variation there is a significant cost associated with sexual fusion and mixing. With rare fusions, selfish proto-mitochondria arise and stay confined within distinct lineages, and are easily eliminated by selection, which results in high mean fitness. Increasing fusion rate allows for a limited spread of deleterious symbionts, leaving the rest of the population mutant-free. A fast transition occurs after the critical rate of cytoplasmic mixing is reached (b), at which point most of the population is overtaken by the selfish proto-mitochondria. Endosymbiont mutation rate is set to 10^−5^.

**Figure 5.**
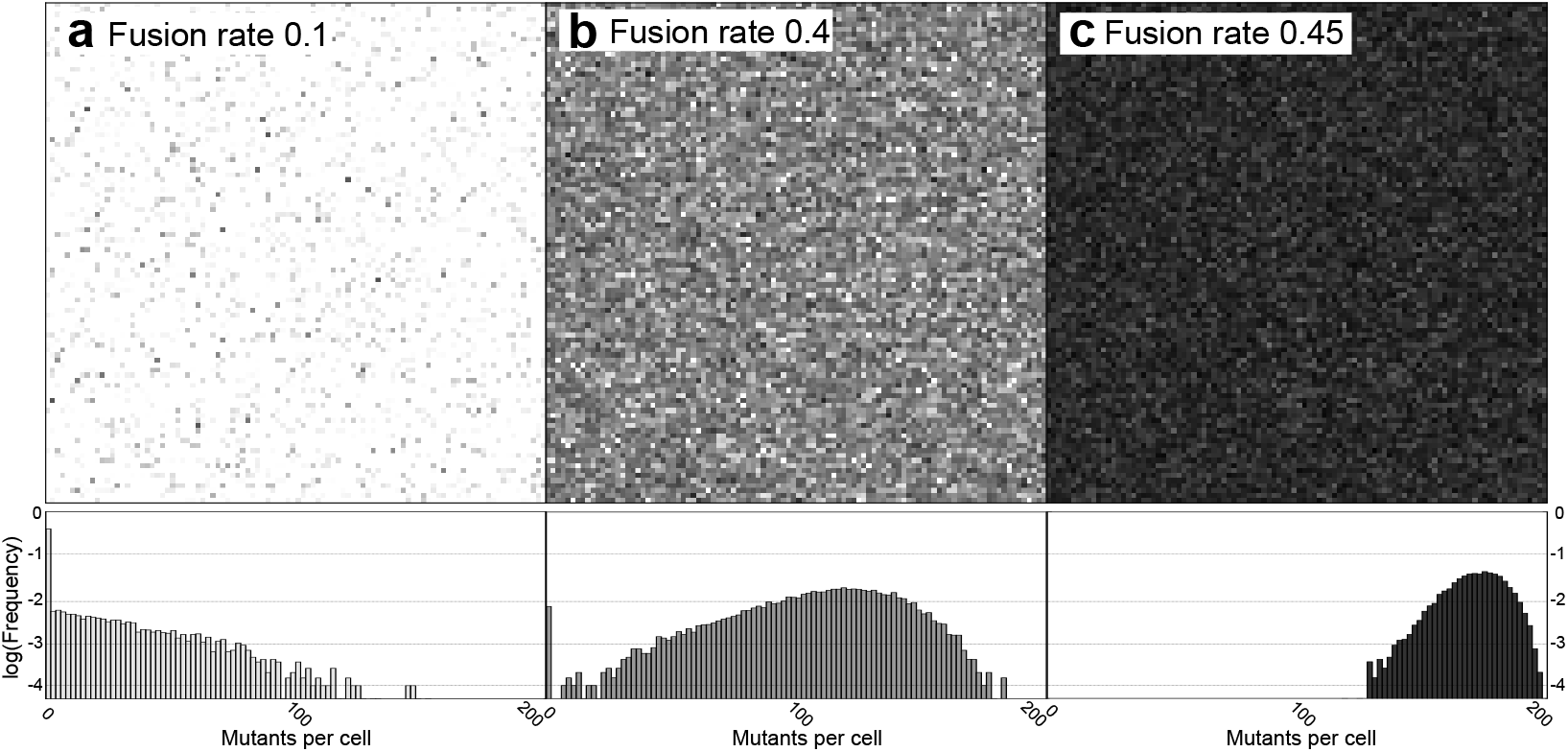
Schematic representation of the population in three equilibrium states with individual pixels representing one of 10000 individuals each. The shade corresponds to the number of mutants *m*. The number of endosymbionts per cell is *M*=200, selfish advantage of the mutants is set to *k*=0.2 (see Figure 4b for the mean population fitness). Note the sudden proliferation of mutants between the panels b and c corresponding to the fast transition to the low-fitness state in Figure 4b. As we demonstrate in the histograms below, the transition is mostly due to reduced variation and extinction of the mutant-free part of the population.

Increasing the frequency of fusions between higher-level units allows selfish symbionts to spread horizontally and “dilutes” the cytoplasmic contents. Similarly to the previous findings in the context of uniparental inheritance of mitochondria [41, 42], we find that mixing reduces inter-individual variation in fitness, at the same time reducing the efficacy of selection on the higher level. As a consequence, the number of selfish endosymbionts grows rapidly as whole-cell fusions become more frequent. The spread of mutants is also favoured by increasing proto-mitochondrial population size within the cytosol, as the random partitioning of symbionts between the daughter cells generates less fitness variation with the higher number of segregating units *M* [45]. On the other hand, small number of symbionts per host cell has an opposite effect and supports higher levels of cooperation (for *M*=20 per cell see Fig. SI1).

Interestingly, in this type of dynamic equilibrium the population fitness might increase with higher mutant reproductive advantage *k* (Figure 4b). Remember, that mutant endosymbionts arise only occasionally with probabilities of only 10^−5^ – 10^−4^, and increase in number mostly due to higher replication rate. As the low fusion rate prevents the horizontal spread of mutants, high *k* results in rapid elimination of higher-level units in isolated lineages dominated by selfish endosymbionts, increasing the mean population fitness.

Finally, with frequent cytoplasmic mixing, population fitness drops to its minimal value, which might involve a sudden step-like transition (Figure 4b). The transition occurs when the rate of mixing becomes high enough to eliminate the mutant-free part of the population (Figure 5), which significantly reduces the inter-individual variance in fitness.

### Spread of cell fusion encoded by the higher-level genome

Now consider an evolutionary scenario where the reproduction mode of the higher-level unit is controlled by a single locus in the haploid proto-nuclear genome, *a*/*A*. The modifier *A* causes the host cell to temporarily fuse with a randomly selected partner, which does not necessarily have to be a carrier of the same allele. We start with an ancestral population of clonally reproducing individuals fixed in *a*, and after equilibration introduce the mutant allele *A* at the frequency of 0.05.

Many eukaryotic species are facultatively sexual and engage in fusion and recombination only under certain environmental conditions, e.g. starvation or oxidative stress. Several rounds of clonal reproduction in-between sexual generations alters fitness variation between higher-level units, raising the possibility that modifiers coding for facultative fusion might evolve easier than with frequent sex. To account for the effect of sporadic cell fusion we introduce a parameter *r*, i.e. probability that *A*-type host successfully induces fusion with a random partner within a single generation.

Due to genetic drift and stochastic fluctuations in finite populations, measuring the modifier frequency is unpractical. Of particular importance in this case is the probability that a mutant allele *A* overtakes the population by completely replacing the wild-type version of the same gene *a—*the fixation probability that can be used to determine whether an allele is favoured or opposed by selection. For a neutral mutant evolving via genetic drift alone the fixation probability equals its initial frequency *p*_0_=0.05, as after a large number of generations the whole population must consist solely of the descendants of a single individual at the generation 0. The fixation probability exceeds *p*_0_ if *A* is favoured by selection, and is lower than *p*_0_ if selection opposes its spread. In practice the fixation probability is calculated numerically, by repeatedly running the simulation until either extinction or fixation of *A* is reached.

Despite the long-term fitness cost brought into the population by *A* (Figure 4), the spread of the allele can be favoured by selection, but only for low values of replicative advantage *k* (Figure 6) and only in the case of a concave fitness relationship (see *Supplementary Information* section B). Though somewhat counterintuitive, the evolution towards sex and lower mean population fitness can easily be explained in terms of the leakage of fitness benefits and the properties of the fitness function used in our modelling [42].

**Figure 6.**
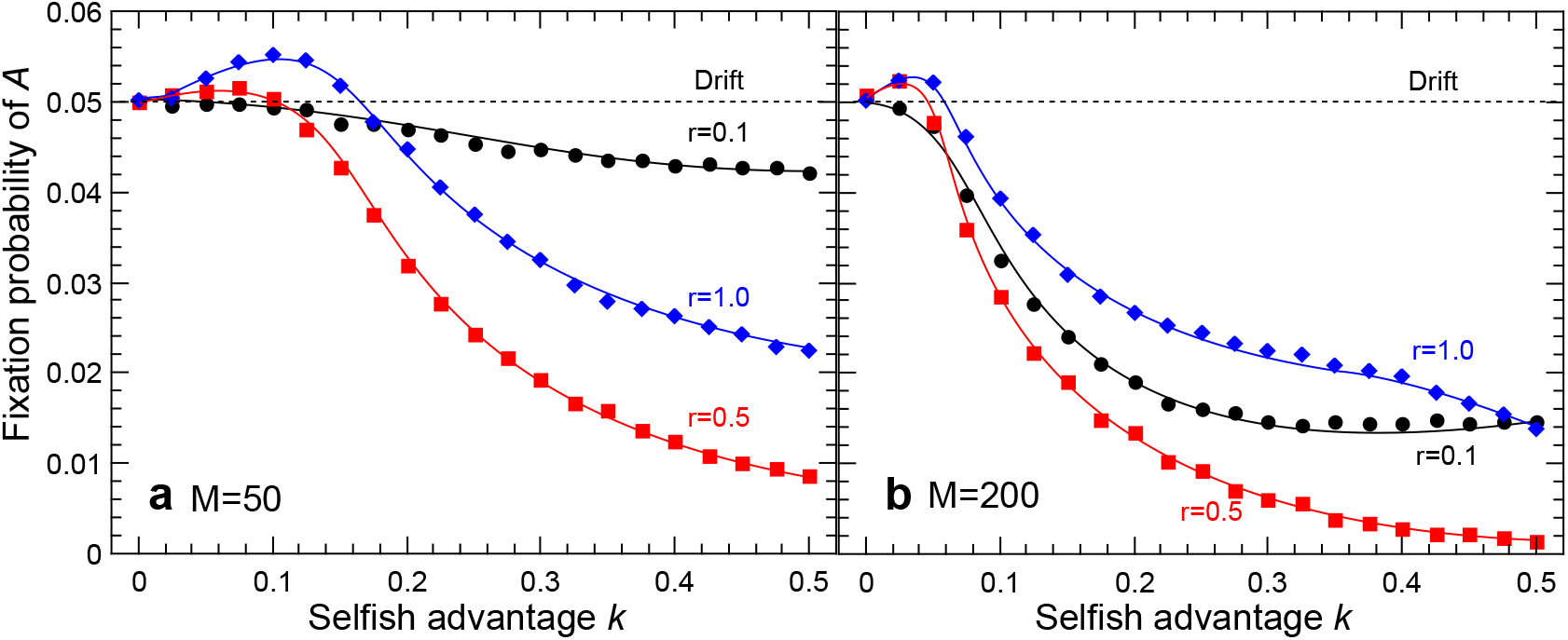
Fixation probability of fusion-inducing allele *A* invading an ancestral population of clonal individuals as a function of mutant replicative advantage *k* for three values of fusion frequency *r*. The number of endosymbionts per cell is 50 (a) or 200 (b). Neutral mutants fix with probability 0.05 (dashed lines). Endosymbiont mutation rate is set to 10^−4^.

Clonal reproduction maintains high inter-individual variation in proto-mitochondrial mutation load, while cytoplasmic mixing generates more individuals with average numbers of mutant endosymbionts. Reduced variation in fitness of the higher-level units results in less efficient selection and a long-term disadvantage. When rare, the carriers of the invader allele *A* fuse almost exclusively with *a*-individuals, thus inheriting many of the cooperative symbionts from the subpopulation of clonal higher-level units. Due to the concave shape of the fitness function, there is a short-term individual advantage to cytoplasmic mixing, as on the average reduced variation removes more low fitness individuals than high. As mating partners are chosen randomly every generation, invader allele *A* manages to maintain its fitness advantage over the clonal subpopulation, even though its invasion is deleterious to the population as a whole. The advantage disappears in the modified case with linearly decreasing fitness function (see *Supplementary Information*).

The fusion-coding allele *A* is favoured by selection only for low values of replicative advantage *k* and high *r*, however. In populations where selfish proto-mitochondria proliferate significantly faster than their cooperative counterparts, sexual mixing becomes strictly disadvantageous. Increasing mutant load strongly opposes the spread of *A,* and so does the increasing endosymbiont population size *M.*

Strikingly, the lower frequency of *A*-coded fusions (*r*<1) does not make the allele more advantageous—the parameter range where *A* is favoured over neutral mutations shrinks with less frequent sex. As *r* decreases, the short-term advantage of reduced variation (which requires constant mixing with random, potentially cooperative partners) is lost making low variation deleterious and more visible to selection. The disadvantage of cytoplasmic mixing loses its significance only when *r* approaches zero and *A* becomes essentially neutral (Figure 6).

Taken together these results imply that sex at the origin of eukaryotes may have had an immediate cost associated with the competition within communities on the lower level of individuality. The disadvantage of sexual proto-eukaryotic fusions is greatest with intense competition and large populations of lower-level units. These findings suggest that putative hosts using some early form of sexual reproduction before acquiring bacterial endosymbionts had little chance of maintaining stable symbiosis and proceeding with the evolutionary transition towards complex eukaryotes.

If the assumptions of our model are correct, an archaeal host to the proto – mitochondria must have reproduced clonally for some time following the initial association. Stable endosymbiosis involving the suppression of selfish reproduction on the lower level (lowering *k*) had to be established before it was safe enough to engage in sexual reproduction with cytoplasmic mixing. Assuming the linear or convex fitness association between the levels of selection, sex is never evolutionary advantageous even with conflicts suppressed, as in these cases reduced inter-individual variation is always costly. Our results therefore firmly support the view of sex as one of the eukaryotic features not present in the earliest stages of eukaryogenesis, and appearing only before the emergence of the true nucleus.

## Discussion

Considerable progress has been made in understanding the endosymbiosis that gave rise to mitochondria [2 – 5]. Nevertheless, the process of eukaryogenesis remains opaque [6]. Insight into this process can be gained to the degree that definitive eukaryotic features can be considered as consequences to the endosymbiosis (e.g., the nucleus [7]). Considerations of the sequelae to endosymbiosis can be broadened to a general evolutionary analysis of conflict and cooperation. As with other levels-of-selection transitions, eukaryogenesis required that conflicts among the lower-level units (the host and the proto-mitochondria) be mediated in order for the higher-level unit (the eukaryote) to emerge.

Initial conflicts resulted primarily from lower-level units competing within the cytosolic environment. Damage within the shared cytosol and selective digestion of lower-level units favoured metabolic regulation and homeostasis. A number of features of eukaryotic metabolism may have evolved in this context, as exemplified by signalling with calcium [27], cAMP [26], and other molecules [25]. At least some of these features were likely already present in the host or symbiont and had only to be recruited into new functions related to conflict mediation. Later in the process of eukaryogenesis, specialized mitochondrial carrier proteins evolved in service of metabolic regulation as exemplified by the adenine nucleotide translocase and membrane uncouplers. The effects were powerful but straightforward. Eukaryotic signalling in general and mitochondrial involvement in particular may originally derive from mechanisms of conflict mediation that maintained metabolic homeostasis.

Informational systems of eukaryotes also bear the imprint of conflict. Early bombardment of the host genome by damaged symbiont DNA drove the formation and growth of the proto-nucleus, but at the same time increased the mutation rate and destabilized the genome. Later in the transition, eukaryotic innovations introduced new cycles of conflicts that required mediation.

Foremost among these innovations was eukaryotic sex. By allowing horizontal transmission of the lower-level units and mixing of symbiont populations, sex counter-intuitively reduced fitness variation between higher-level units and in this way was potentially disruptive and had extensive ramifications. We showed that fitness costs due to competition between proto-mitochondria within the shared cytosol increase with more frequent cell-cell fusions and larger endosymbiont populations. Although the ability of ancient archaeal hosts to fuse could have facilitated the uptake of endosymbionts, whole cell fusion was likely to destabilise the newly emerging community and lead to failure of the evolutionary transition. The archaeal host was better off reproducing clonally for at least some time following the initial endosymbiotic event. A stable community involving suppressed competition on the lower level had to be established before it was safe enough to engage in sexual reproduction and cytoplasmic mixing.

Traditionally the evolution of sex has been analysed in the context of recombination among nuclear genes, DNA repair and associated fitness of the eukaryote. It is not clear, however, whether the same evolutionary forces were responsible for the origin of between-host fusion during the eukaryogenesis, when the higher-level unit was still emerging. It has been suggested before that the evolution of sex in the form of temporary cell fusion could have been promoted by endosymbionts producing oxygen free radicals under the conditions of low metabolic demand, e.g. if the host was unable to grow due to mutations in the proto-nuclear genome and therefore had little ADP to offer [16]. ROS is known to induce the formation of gametes and sex in some algae today [46]; it would not be surprising if the connection between ROS and sex predated the eukaryogenesis itself. It is well established, for example, that DNA damage generated by UV radiation and possibly ROS serves as a trigger for species-specific cellular aggregation and DNA repair via recombination in Archaea [47, 48] and, similarly, can increase the rate of transformation in bacteria [49]. As a side effect of ROS-induced cell fusion, proto-eukaryotes therefore had an opportunity for regular recombination and repair of their unstable chromosomes, or were simply able to mask mutations with clean copies of the same genes, restoring the growth and favourable conditions for the endosymbiont proliferation.

Cell fusion, recombination and genome repair were critical for the formation of large eukaryotic genomes, given that intense horizontal symbiont gene transfer and intron bombardment was taking place [7, 8, 39]. From this point of view, the origin of sex as proto-eukaryotic fusion was a prerequisite for the evolution of stable complex cells with large nuclear genomes. Given the results of our modelling, we therefore argue that sex represents a late proto-eukaryotic innovation, allowing for the growth of the chimeric nucleus and contributing to the successful completion of the evolutionary transition.

Evolutionary analysis of competition and cooperation restricts the order of key events leading to the emergence of the eukaryotic cell. Large stable nuclear genome—the key feature of the group—required repair and recombination across the full length of the chromosomes and therefore could have been maintained only after the origin of whole-cell fusion. Sex, on the other hand, could not have evolved before the competition on the lower level was suppressed. Similarly, the transfer of mitochondrial genes to the chimeric nucleus, largely contributing to its growth, required relaxed selection on the lower level. The founding members of the mitochondrial carrier family may have originated as proto-mitochondrial innovations, in which case they did not require the presence of mitochondrial protein import machinery, and could have evolved before sex or genome transfer. ATP/ADP translocase and membrane uncouplers, together with the generic signaling pathways recruited from the prokaryotic ancestors, maintained endosymbiont homeostasis and contributed to the suppression of such conflicts on the lower level.

### Data accessibility

The C++ source code for the simulation routines is available at ucl.ac.uk/∼ucbprad/fusionconflicts.

### Competing interests

The authors have no competing interests related to this work.

### Authors’ contributions

AR developed the model and carried out the simulations. AR and NWB interpreted the simulations and wrote the manuscript.

## Acknowledgements

The authors acknowledge the use of the UCL Legion High Performance Computing Facility (Legion@UCL), and associated support services, in the completion of this work. Zena Hadjivasiliou, Nick Lane and Andrew Pomiankowski provided helpful comments.

## Funding statement

Radzvilavicius conducted this research as a recipient of UCL’s Bogue Fellowship Award with additional funding from EPSRC/CoMPLEX PhD studentship.

## SUPPORTING INFORMATION TO

### A. Small number of endosymbionts

**Figure S1.**
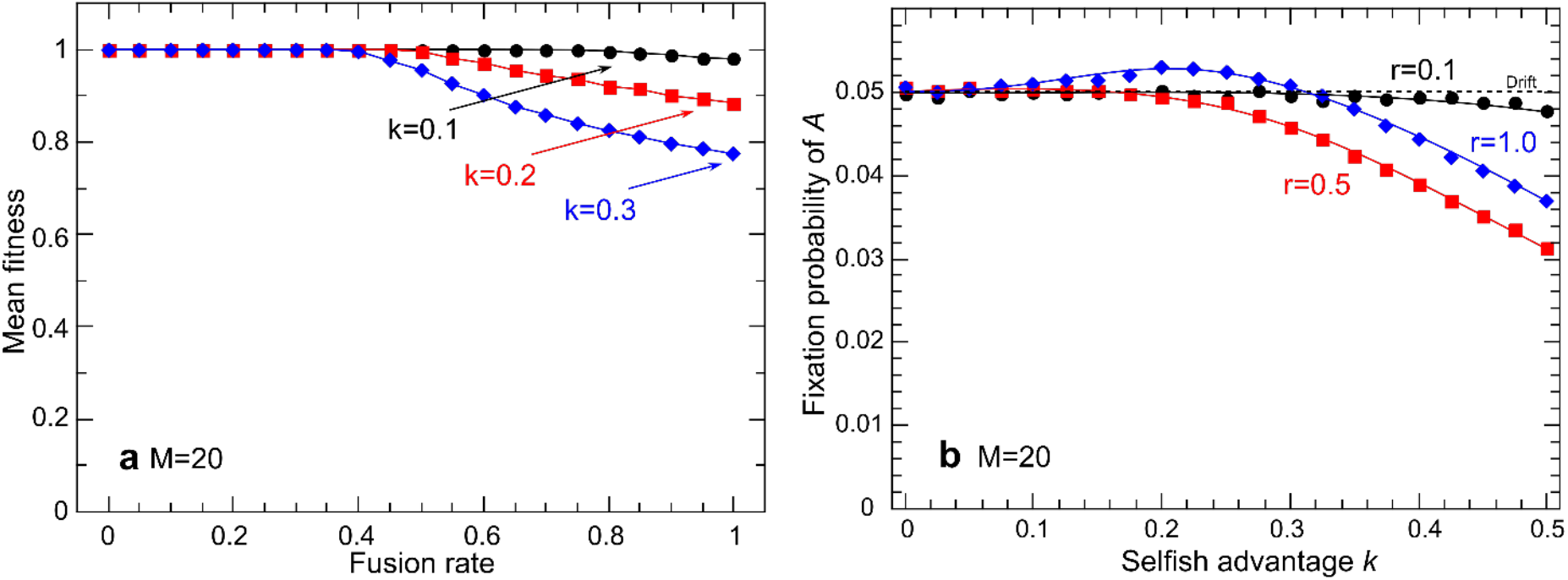
(a) Mean population fitness as a function of the fusion rate for three values of selfish replication advantage *k* with M=20 endosymbionts per cell. The loss of fitness is less significant compared to the cases with M=50 and M=200 endosymbionts per cell. (b) Fixation probability of the allele *A,* coding for the whole cell fusion invading the ancestral population of clonal individuals with M=20 endosymbionts per cell. Neutral mutant fixes with the probability of 0.05. *r* represents the fusion rate of the proto-sexual individuals bearing the allele *A*. Cell fusion is evolutionary advantageous for a wider range of *k*, but only if the cell-cell fusion events are frequent (*r*=1 in this case).

### B. Linear and convex fitness functions

In the main text we assumed that the fitness association between the contents of the cytoplasm and the higher level unit follows a concave function 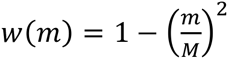(Fig. S2). This particular assumption, however, is based on the contemporary association between mitochondrial function and the fitness of the eukaryotic cell. It could therefore be informative to consider alternative relationships of the form 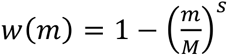, as the shape of the fitness function is directly responsible for the strength of selection on the higher level, and alters the way increased variation affects the fitness of an individual. Here we restrict ourselves to the linear (*s*=1) and convex (*s*=0.5) fitness associations as shown in Figure S2. The case of a convex fitness function is hard to justify biologically, as it implies that fitness declines fast with few mutant proto-mitochondria, and flattens once the higher level unit is gradually overtaken by selfish endosymbionts. This seems to be against both intuition and empirical evidence but is included here for the sake of completeness.

**Figure S2.**
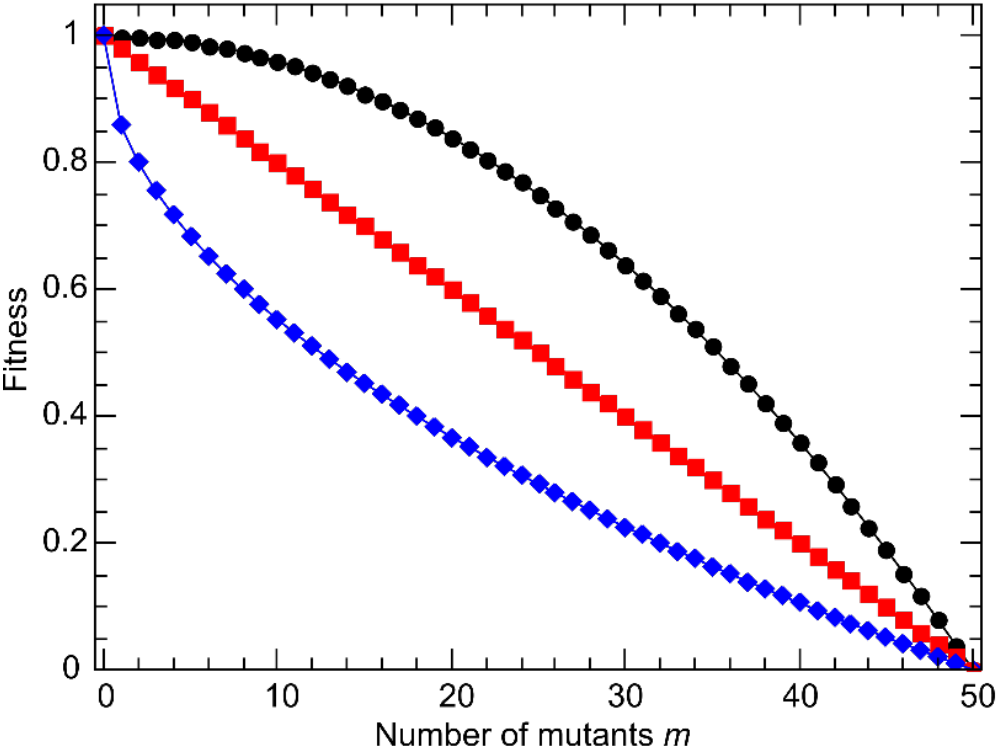
Association between the fitness on the higher level and the number of deleterious endosymbionts *m*: concave (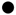, s=2), linear (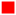, s=1) and convex (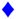, s=0.5) fitness functions.

For the linear fitness association, we find that higher rate of cell-cell fusion is needed to cause a significant decrease in fitness (Fig. S3, a), but once it is reached the mean population fitness rapidly declines with both fusion rate and reproductive advantage of mutants *k*. Cytoplasmic mixing and lower variation between individuals on the higher level does not have an immediate short-term advantage as it did in the concave case. As a consequence, cell-cell fusion is never evolutionary advantageous, as demonstrated by the fixation probability of a fusion-causing allele *A*, which is in all cases lower than the corresponding probability for the neutral allele (Fig. S4, a).

With the convex fitness function, small initial changes in the number of mutants *m* cause an immediate decline in higher-level fitness. Lineages in which selfish mutants first arise are therefore eliminated rapidly by the natural selection, which is reflected in high and constant mean population fitness independent of both *k* and fusion rate (Fig. S3, b). As the overall number of mutant endosymbionts within the population is kept low at all times, the cellular fusion is never advantageous (Fig. S4, b); the fixation probability of the fusion modifier *A* stays close to the neutral case for most parameter values.

**Figure S3.**
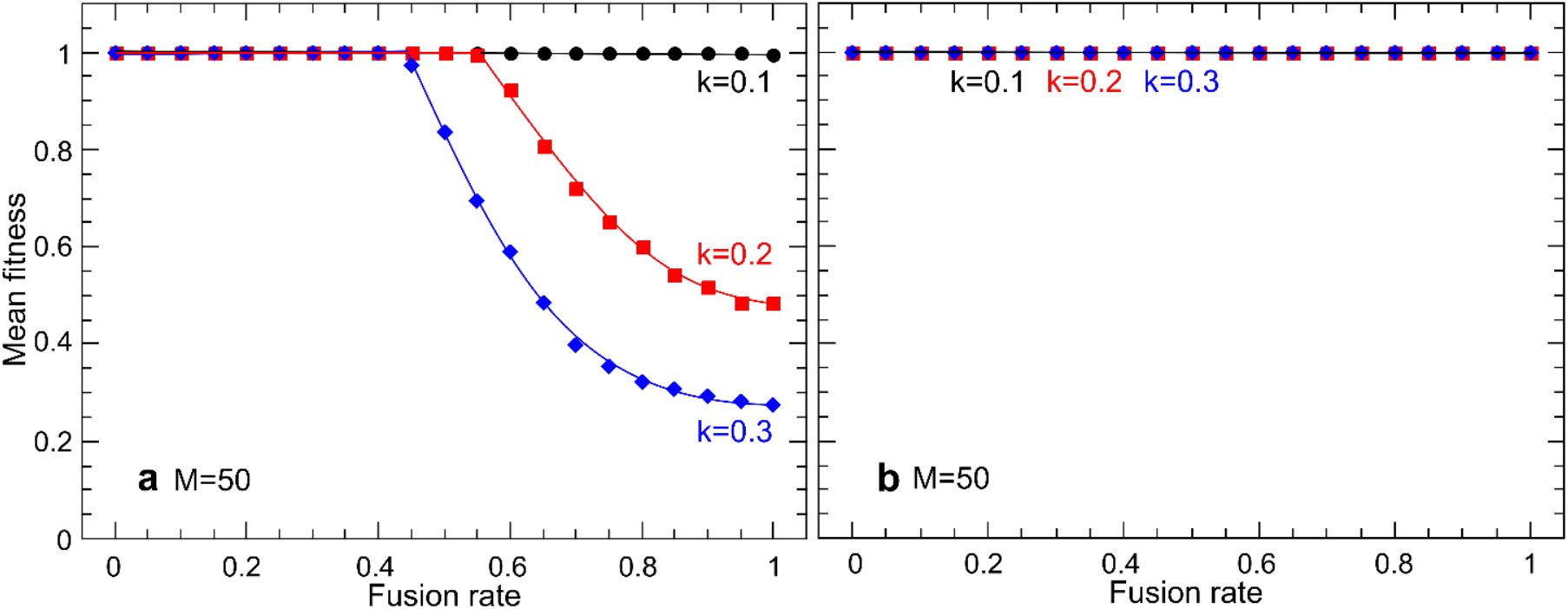
Mean population fitness as a function the inter-cellular fusion rate for linear (a) and convex (b) fitness associations. The number of proto-mitochondrial endosymbionts is *M*=50.

**Figure S4.**
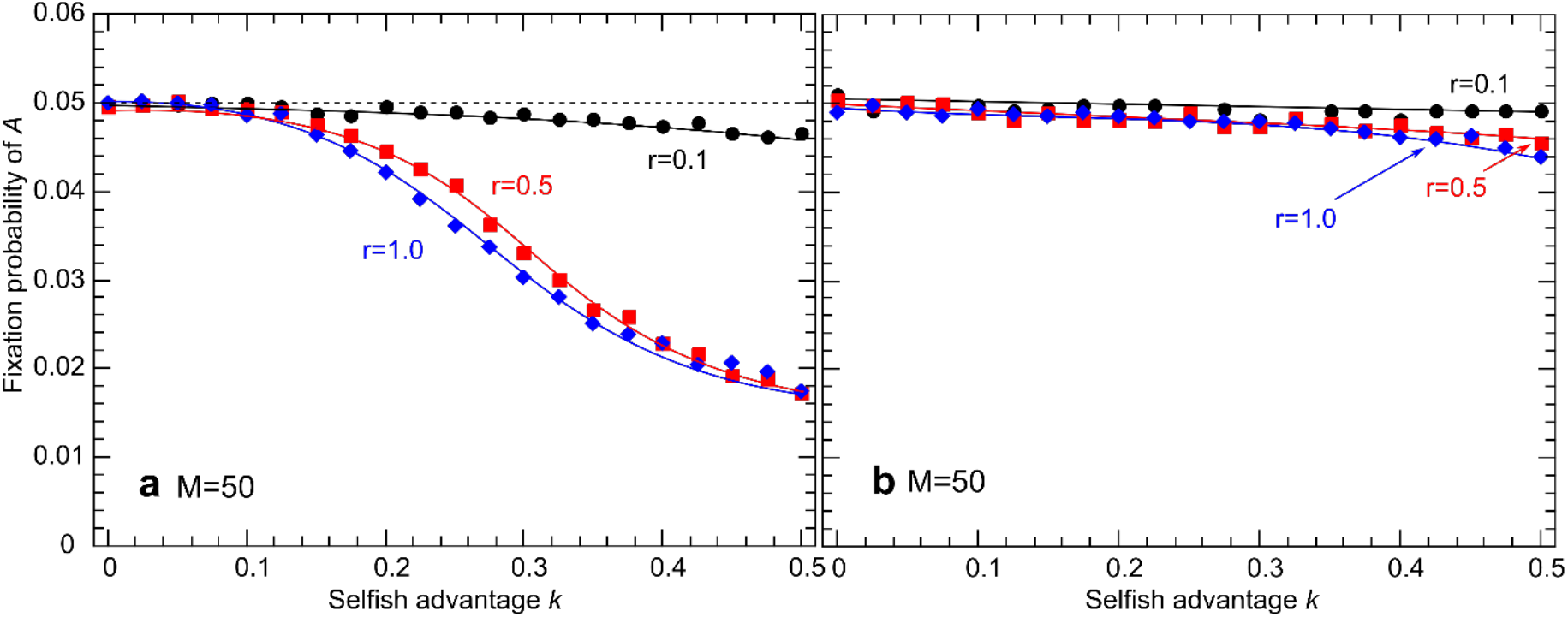
Fixation probability of the fusion-inducing allele *A* as a function of the selfish advantage of mutant endosymbionts *k* for linear (a) and convex (b) fitness associations. Neutral mutant fixes with the probability of 0.05 (dashed line). *r* represents the fusion rate of the proto-sexual individuals bearing the allele *A*.

### C. Stochastic processes in the model life cycle

Symbiont segregation between two daughter cells during clonal division was modelled as a simple random sampling without replacement. Probability to give birth to a daughter cell with *r* ∈ {0,1,…,*M*} mutant symbionts given that the parental cell had *m* mutants is given by

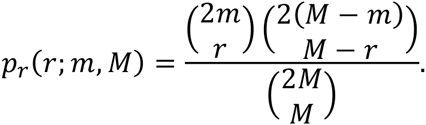

Mutation events on the lower level were assumed to follow binomial probability distribution. Probability for a higher level unit to accumulate *l* new mutants within a single generation is therefore

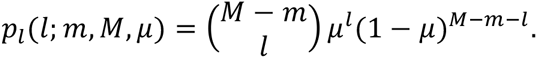

Selection on the lower level was implemented as a single weighted resampling with replacement. Given replicative advantage of selfish symbionts 1 + *k* and *m* selfish symbionts within a higher level unit, the number of defectors after selection *q* follows

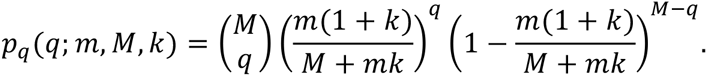

Similarly, selection on the higher level was implemented as sampling with replacement with selection probabilities proportional to fitness values. Random numbers from relevant probability density functions in the C++ implementation of the model were generated by employing “Mersenne Twister” random number generator from GNU Scientific Library.

